# Serum proteomics reveals distinct phenotypic signatures to IL-6 blockade between two immunotherapies

**DOI:** 10.64898/2026.03.27.712461

**Authors:** Catherine Sniezek, Deanna Plubell, Katarina Vlajic, Andy Hoofnagle, Christine C Wu, Jane H Buckner, Devin Schweppe, Cate Speake, Michael J. MacCoss

**Author notes:** Correspondences should be addressed to: Michael J. MacCoss.

## Abstract

A recent clinical study tested the effects of two different monoclonal antibodies (mAbs) (siltuximab, anti-IL6; tocilizumab, anti-IL6R) on the fate and function of T-cells in people with type 1 diabetes. While both mAbs affect the response of T-cells to stimulation, they have very different, sometimes opposing mechanisms. Here, we use mass-spectrometry based proteomics to analyze longitudinal serum samples (baseline and two weeks post-treatment) from 20 clinical trial participants to examine the effects of siltuximab and tocilizumab on extracellular vesicles. To accomplish this, serum samples were enriched for extracellular vesicles with Mag-Net and analyzed by LC-MS/MS to identify significantly differentially abundant protein groups and pathways. Proteome analysis confirmed highly reproducible measurements across multiple draw dates. In total, we quantified >3300 protein groups of which 46 protein groups had significantly altered abundance after mAb treatment. Tocilizumab altered pathways associated with proteostasis (neddylation) and pre-notch transcription and translation. Siltuximab altered FCGR activation pathway members. In addition, quantitation of the monoclonal antibody therapies themselves enabled the measurement of the correlation between drug amounts and impacted proteins. Taken together, this work demonstrates the utility of the Mag-Net method to evaluate the impacts of therapeutic interventions on serum extracellular vesicles.

## Introduction

The measurement of therapeutic response remains one of the most critical challenges in precision medicine. Individual responses to drug treatments vary dramatically based on complex interactions between genetic background, environmental factors, microbiome composition, disease heterogeneity, and drug pharmacokinetics. This variability underscores the need for robust, quantitative methods to monitor treatment efficacy and distinguish responders from non-responders early in therapeutic intervention. Such capabilities are essential not only for optimizing individual patient care but also for understanding the mechanistic basis of drug action and identifying biomarkers that predict treatment success.

Blood-based proteomics has emerged as a powerful approach for monitoring therapeutic response. Plasma and serum are ideal biological matrices for longitudinal monitoring as they can be obtained through minimally invasive procedures, collected repeatedly over time, and contain proteins and extracellular vesicles that reflect the physiological state of virtually all organ systems. Circulating extracellular vesicles integrate signals from diverse tissues, making them a rich source of information about systemic responses to therapeutic interventions^1^. Since serum and plasma are commonly analyzed in clinical laboratories, the measurement of extracellular vesicles can be easily translated from proteomic discovery into clinical practice.

Recent technological advances have dramatically expanded our ability to probe the plasma proteome. The development of mass spectrometers capable of collecting tandem mass spectrometry (MS/MS) data at >100 spectra/second^2–5^, improved sample preparation methods^6,7^, and novel computational approaches^8–10^, now enable the quantification of over 4,000 proteins from individual plasma samples. Particularly important has been the development of enrichment strategies that address the fundamental challenge of plasma proteomics: the enormous dynamic range spanning over 10 orders of magnitude in protein concentration^11^. Extracellular vesicle (EV) enrichment methods have proven especially valuable for plasma proteomics, as EVs contain cellular proteins that would otherwise be masked by highly abundant plasma proteins like albumin and lipoproteins. The Mag-Net method^6^, which uses magnetic bead-based enrichment to isolate EVs through combined electrostatic and size-capture physiochemical properties, has shown particular promise in reducing sample dynamic range while preserving quantitative accuracy.

IL-6 is an important cytokine for diverse cellular processes, from innate and adaptive immunity to cellular differentiation and metabolic control^12^. Dysregulation of IL-6 is implicated in various pathologies, such as inflammation, autoimmunity, and cancer^13^. Two strategies for therapeutic blockade of IL-6 signaling include direct neutralization of the IL-6 cytokine and blockade of the IL-6 receptor. Siltuximab, a chimeric monoclonal antibody, directly binds and neutralizes IL-6, preventing its interaction with both membrane-bound and soluble IL-6R. In contrast, tocilizumab, a humanized monoclonal antibody, targets IL-6R itself, blocking both classical and trans-signaling pathways^13,14^. While both approaches interrupt IL-6 signaling, they may have distinct biological consequences due to differences in their mechanism of action, effects on cytokine clearance, and impact on receptor dynamics.

Our recent mechanistic clinical study provided an opportunity to directly compare these two IL-6 blocking strategies in the setting of type 1 diabetes (T1D)^15^. This study administered either monoclonal antibodies (mAb) siltuximab or tocilizumab to people with T1D and evaluated effects on T cell fate and function. Surprisingly, despite both mAbs targeting the same signaling pathway, the study revealed different, sometimes opposing effects on T cell responses. Among other changes, siltuximab improved the ability of effector T cells to respond to regulatory T cell suppression; this was not observed with tocilizumab. These divergent immunological outcomes raised fundamental questions about how ostensibly similar therapeutic interventions can produce such different biological responses.

The unexpected divergence in cellular immune responses between siltuximab and tocilizumab treatment highlighted the need for a broader systems-level understanding of how these therapies impact the immune system. While previous investigations focused on targeted immune cell populations, the systemic effects of these interventions on the circulating proteome and extracellular vesicles remained unexplored. Proteomic analysis of longitudinal serum samples offers a complementary perspective that can reveal both direct effects of IL-6 pathway blockade and secondary consequences throughout interconnected biological networks.

Here, we present an analysis of extracellular vesicles from serum samples collected during this clinical study, comparing baseline and two-week post-treatment timepoints for both siltuximab and tocilizumab treatments. Our objectives were to: (1) establish the feasibility of using Mag-Net EV enrichment for longitudinal serum samples in people receiving therapies, (2) characterize the global proteomic changes induced by each therapeutic intervention, (3) identify shared and unique protein signatures between the two IL-6 blocking strategies, and (4) relate proteomic changes to the underlying mechanisms of drug action. By leveraging advances in sample preparation, mass spectrometry, and computational analysis, we aimed to provide new insights into how different approaches to blocking the same signaling pathway can produce divergent biological outcomes, with implications for understanding therapeutic mechanisms and optimizing treatment selection in precision medicine.

## Methods

### Sample acquisition

Serum samples from siltuximab (anti IL-6) and tocilizumab (anti-IL-6R) clinical studies were provided as previously described^15^. Each study was single-arm, single-dose, and open label, with each participant serving as their own baseline control. 10 participants from each study provided baseline and 2-week post-treatment serum samples (40 total samples). 5 of these participants were enrolled in both studies, with a minimum of 5 months between studies. All participants (adults, aged 18-45) had type-1 diabetes and were between 4 months to 10 years post-diagnosis. All had at least one diabetes-related autoantibody and had detectable insulin secretion on a mixed-meal tolerance test within 60 days of enrollment. Exclusionary criteria included abnormal complete blood counts or active viral infections. Siltuximab was administered at 11 mg/kg and tocilizumab at 8 mg/kg according to Benaroya Research Institute IRB protocols (siltuximab, IRB15085; tocilizumab, IRB15159). The siltuximab study was registered under an FDA Investigational New Drug application, with consequent registration on ClinicalTrials.gov (NCT02641522). All samples were assayed in a blinded and randomized manner.

### Mag-Net enrichment method

To enrich serum samples for extracellular vesicles, the Mag-Net method was performed as described^6^ with the use of a Kingfisher Flex (ThermoFisher). For each sample, 25 uL of MagResyn SAX beads (Resyn Biosciences) at the supplied 20 mg/mL was equilibrated with 500 uL of bis-tris-propane buffer (100 mM bis tris propane, pH 6.3, 150 mM NaCl). Equilibrated beads were applied to 100 uL of serum or reference plasma treated with HALT protease and phosphatase inhibitor (Thermo Fisher Scientific). Protein-captured beads were washed three times with bis-tris-propane buffer and then reduced with 10 mM TCEP in 50 mM Tris, pH 8.5, and 1% SDS for 60 minutes at 37 C. 800 ng of enolase was added to each sample for an internal process control. Reduced proteins with capture beads were alkylated with 15 mM iodoacetamide for 30 minutes, then quenched with 10 mM DTT. Reduced and alkylated proteins with capture beads were precipitated with 70% acetonitrile, washed three times with 95% acetonitrile, then twice with 70% ethanol. Precipitated proteins were digested with trypsin (7.5 ug per sample) and incubated at 47 C for 1 hour. Peptides were quenched with 0.5% trifluoroacetic acid, mixed with Pierce Retention Time Calibrant peptide cocktail to 50 nmol/uL (Thermo Fisher Scientific) and stored at -80.

### Mass Spectrometry

All samples were analyzed with a Thermo Scientific Orbitrap Astral coupled to a Vanquish Neo UHPLC and Easy Spray Source. All LC runs had a flow rate of 1.3 µL/minute and were 24 minutes long with a 4-55% B gradient over 20.5 minutes, followed by a 99% B wash. (Mobile phase A: 0.1% formic acid in water. Mobile phase B: 80% acetonitrile, 0.1% formic acid in water. Column: Pepsep 15 cm by 150 µm, 1.9 µm bead size) Spectra were acquired in DIA mode with 4 Th wide optimized windows^16^. Precursor isolation windows spanned 400-900 Th, and MS1 spectra were acquired in the Orbitrap at a resolution of 240,000. HCD fragmentation mode was used at 27% NCE and MS2 spectra were acquired with a normalized AGC target of 250% and a 375-985 Th scan range.

### Data Processing and Quantification

A custom FASTA database was constructed by appending the amino acid sequences of siltuximab (anti-IL-6) and tocilizumab (anti-IL-6R) to the canonical human proteome (UniProt, Swiss-Prot) supplemented with common contaminant sequences and yeast enolase. Vendor raw files were converted to mzML format using MSConvert (ProteoWizard)^17^.

A fine-tuned spectral library was generated using the search results from DIA-NN v1.8.1 with the Carafe deep learning–based predictor^18^ as part of a Nextflow workflow^19^. A single representative file (Ast_20240220_S10_26.mzML) was searched against the custom FASTA database in “library-free mode” to produce training data for Carafe to fine-tune the AI predicted retention times and fragment intensities. Then all mzML files were then processed with DIA-NN v1.9.2 using the fine-tuned Carafe library with match-between-runs enabled, the unrelated runs option selected, and neural network classifier set to conservative mode.

DIA-NN search results were imported into Skyline (v24.1 via File > Import > Peptide Search using the skyline.speclib spectral library output. Transition settings were configured to include precursor charge states 2+ and 3+, product ion charge states 1+ and 2+, and y, b, and precursor ion types, with product ions ranging from ion 3 to the last ion within an m/z range of 200–2000. Up to 6 product ions were selected per precursor with a minimum of 4 required. Full-scan settings specified centroided data for both MS1 and MS/MS levels with a mass accuracy of 10 ppm each. MS1 filtering included 3 isotope peaks, and MS/MS extraction used DIA acquisition with an isolation scheme derived from the results files. Retention time filtering was applied using a ±3-minute window around MS/MS identifications.

Protein inference was performed in Skyline using parsimony-based protein grouping with the option to find the minimal protein group list that explains all peptides, removal of subset protein groups, shared peptides duplicated between protein groups, and a minimum of 1 peptide per protein group. This yielded a final document containing 3,318 protein groups, 25,816 peptides, 29,968 precursors, and 268,197 transitions. Peptide quantification was normalized by equalized-medians. For peptides not identified in individual replicates, peak integration boundaries were imputed by transferring boundaries from the exemplary replicate using Skyline’s built-in functionality. Raw data and Skyline documents available at Panorama Public (https://panoramaweb.org/IL6-Biologics-Serum.url, ProteomeXchange ID PXD076219).

### Post-hoc data analysis

Quantitative peptide- and protein-level reports were exported from Skyline for downstream statistical analysis. R version 4.3.2 was used to manipulate the peptide data matrix. Owing to their significant sequence homology with endogenous immunoglobulins, therapeutic antibodies (siltuximab and tocilizumab) were quantified using only unique peptides which showed a significant quantitative difference over untreated samples. All other peptides were collapsed to protein groups by summing assigned peptides, and protein quantitation was normalized across samples to equalize medians. The linear modelling tool Limma^20^ was used to determine proteins differentially altered by therapeutic antibody treatment, pairing each pre- and post-treatment sample. P-values were corrected by the Benjimini-Hochberg approach with a significance cutoff of P-adjusted < 0.05. Gene-set enrichment analysis was performed with the fgsea tool with the reactome^21^ or Jensen’s compartments^22^ databases, with minimum set size 5 proteins and maximum size 500. R code available at GitHub (https://github.com/uw-maccosslab/manuscript-Il6-biologics-serum).

## Results

### Mag-Net EV enrichment is reproducible for longitudinal studies

Meaningful analyses of plasma or serum samples from a longitudinal clinical study requires reproducible measurements across the time span of the trial. Sample handling factors such as contamination of plasma samples with erythrocytes have been shown to impact plasma proteomics analyses, especially in bead-based enrichment methods, such as Mag-Net used here ^23^. Therefore, it must be established that Mag-Net enriched serum proteomics is reproducible across the analysis of many samples.

The tocilizumab and siltuximab studies included five individuals who participated in both trials. These five people provided baseline serum samples prior to the siltuximab trial and again a year later prior to enrollment in the tocilizumab trial (Figure 1B). Paired baseline extracellular vesicle proteomes were highly correlated within an individual between participant baseline draws. (Spearman rho > 0.95). Unsupervised clustering of Spearman rho correlation coefficient values indicated that longitudinal proteome profiles from each individual were consistently clustered with one another (Figure 1C). To evaluate sources of variance across all 40 samples from both studies, principal component analysis was performed. Samples were not clustered by their study of origin. Instead, higher variability was observed in the baseline sample group than in the post treatment group, indicating a strongly similar response to treatment across both siltuximab and tocilizumab (f-test p-value = 0.0001) (Figure 1D). These data establish that Mag-Net sample enrichment of serum samples is highly reproducible and robust to sample collection spanning at least 1 year, but maintains sensitivity to systematic changes to the extracellular vesicle proteome.

**Figure 1.**
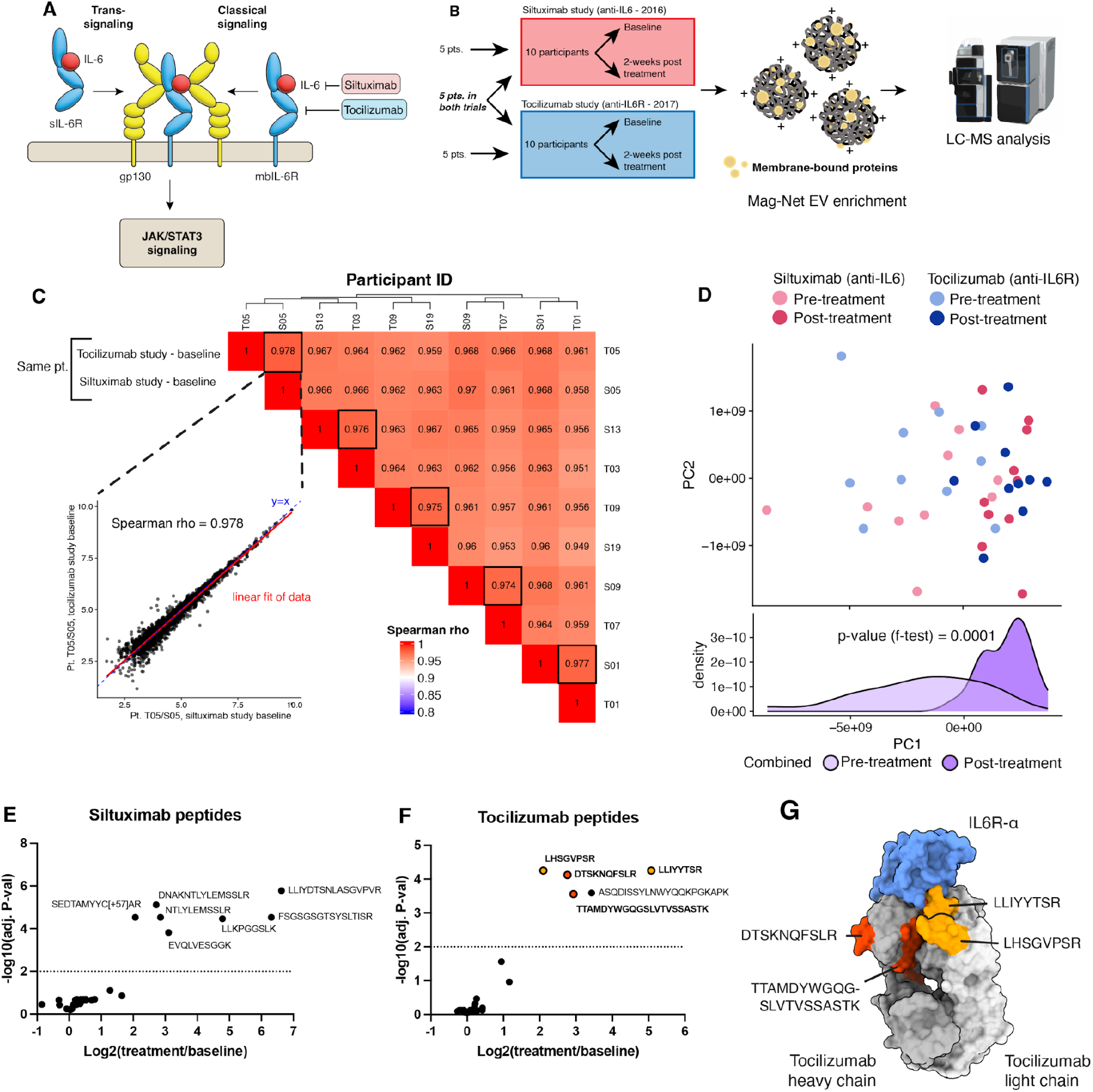
Analysis of Mag-Net derived EV fraction enriched from longitudinal serum samples. (A) Schematic representation of the IL6 pathway with relevant mAb inhibitors (B) Proteomic workflow for the analysis of serum samples from two type 1 diabetes trials. (C) Spearman rho correlation coefficients of baselines from the 5 individuals enrolled in both drug trials. Inset shows correlation of the same individual with blood draws spanning one year. (D) Principal component analysis of baseline and post-treatment samples from both drug trials. (E-F) Peptide-level quantitation of enriched serum-derived siltuximab and tocilizumab peptides. (G) Quantified tocilizumab peptides overlaid with structure of tocilizumab and IL-6R (PDB 8J6F).

### Characteristic peptides from monoclonal antibody treatment are quantified

The serum half-lives of the monoclonal antibody therapies are 20.6 and 11 days, respectively, for siltuximab and tocilizumab (Speake et al. 2022). Thus, we expected that these 14-day serum samples would contain detectable levels of siltuximab and tocilizumab. Bottom-up, serum proteomics quantified 44 peptides (7 unique) matching siltuximab, and 41 peptides (5 unique) matching tocilizumab. However, significant sequence homology between the humanized, monoclonal antibody therapies and other endogenous immunoglobulins obscures the quantitation of these therapies directly when all matched peptides are aggregated to a protein-level quantitation value. To resolve this, peptides matching to siltuximab or tocilizumab were individually examined for significant changes before and after treatment, and only a subset (unique peptides) of siltuximab and tocilizumab peptides were found to change with treatment. (Figure 1, E-F). Further, of the five tocilizumab peptides changing with treatment, four of them are found in the variable domain of the antibody, interfacing with the IL-6R epitope. (Figure 1G) These data demonstrate that a peptide-centric approach can effectively quantify proteins with significant sequence homology with other proteins in the sample matrix.

### Monoclonal antibody therapy induces proteome-wide abundance changes in enriched serum samples

Samples for extracellular vesicle analysis were collected longitudinally, once before treatment and once two weeks after treatment. Because of this, each participant acts as their own control for differential protein enrichment analysis. Analysis by Limma ^20,24^ of paired treatment and baseline samples showed that tocilizumab treatment significantly altered the serum abundance of 42 protein groups and siltuximab treatment altered 9 protein groups. Of these, 5 protein groups were altered by both treatments (ELAP2, CO4A, CO4B, KPSH1, and CCD83). (Figure 2, A-B)

**Figure 2.**
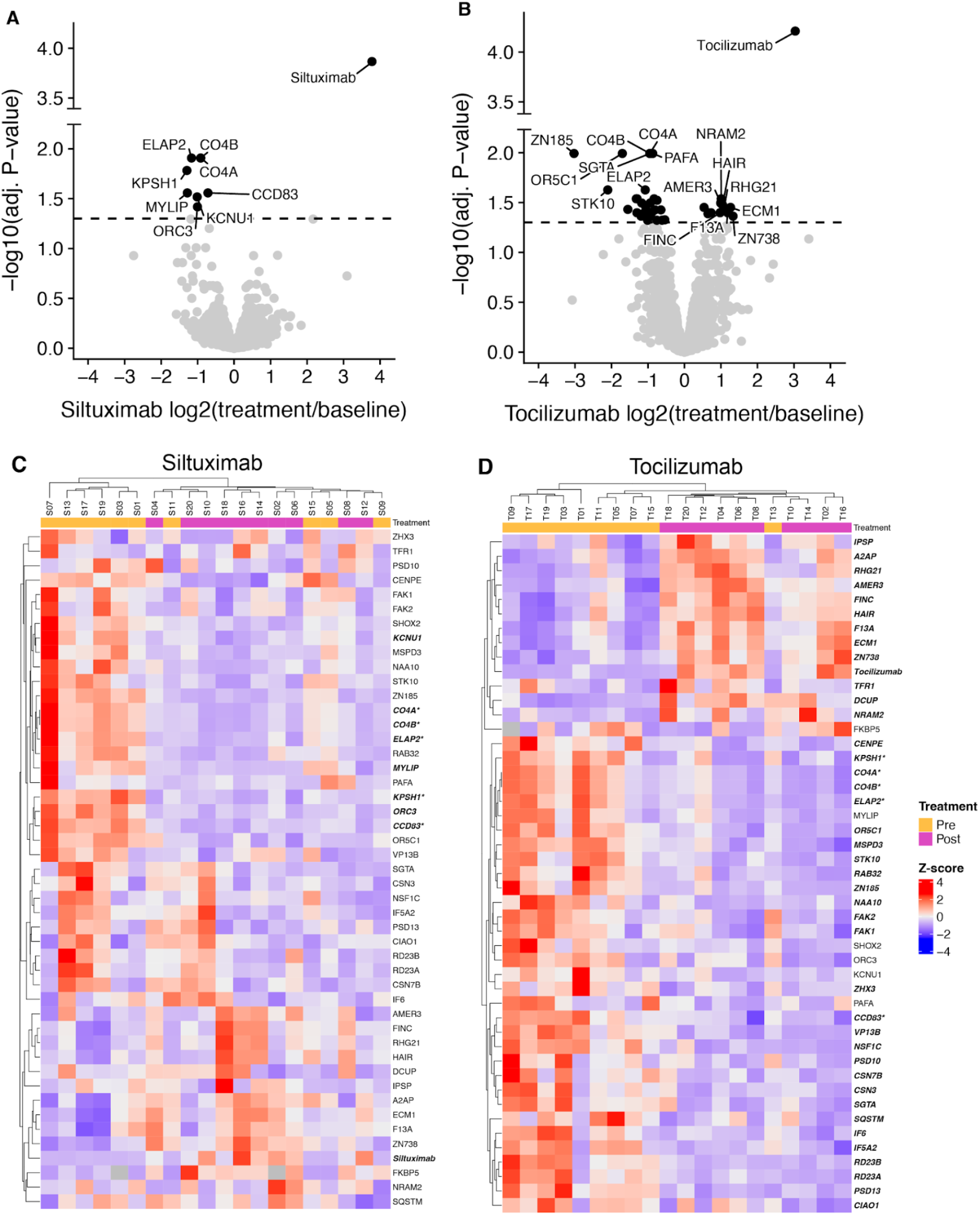
Therapeutic antibody treatments show protein-level impact on serum proteome. (A-B) Volcano plot showing limma results for siltuximab and tocilizumab. P-values corrected by Benajimini-Hochberg approach. (C-D) unsupervised clustering of protein-level, sample-wise Z-scores.

The 46 protein groups found to be altered by either treatment were used for unsupervised clustering of samples within treatments according to their protein-wise z-score. For tocilizumab, all but one sample clustered according to treatment status. (Figure 2D). The siltuximab trial showed less clustering according to these protein groups (Figure 2C). Taken together, these data show that individually, each treatment had a measurable effect on the serum proteome two-weeks after a single administration, and that tocilizumab had a more pronounced impact than siltuximab.

### IL-6 blockade by tocilizumab or siltuximab impact the serum proteome on a pathway level

Once we established that tocilizumab and siltuximab each exhibit significant impacts to the EV-enriched serum proteome, we investigated whether the directionality of these impacts were similar across treatment groups. We find that the effects of each treatment show similar directionality for many proteins, excepting the therapeutic antibodies themselves (Figure 3A), with a Pearson correlation coefficient across the whole set of R = 0.37, and for only proteins significantly impacted by either treatment R = 0.61. Among the proteins altered (decreased) by both treatments is complement factor 4 (Figure 3B), which agrees with previous studies demonstrating that IL-6 blockade by tocilizumab reduces blood-derived concentration of complement factor 4^25^. In addition, tocilizumab more than siltuximab increased serum concentration of fibrinonectin (FINC), which is also a known result of IL-6 blockade (Figure 3B)^12^.

**Figure 3.**
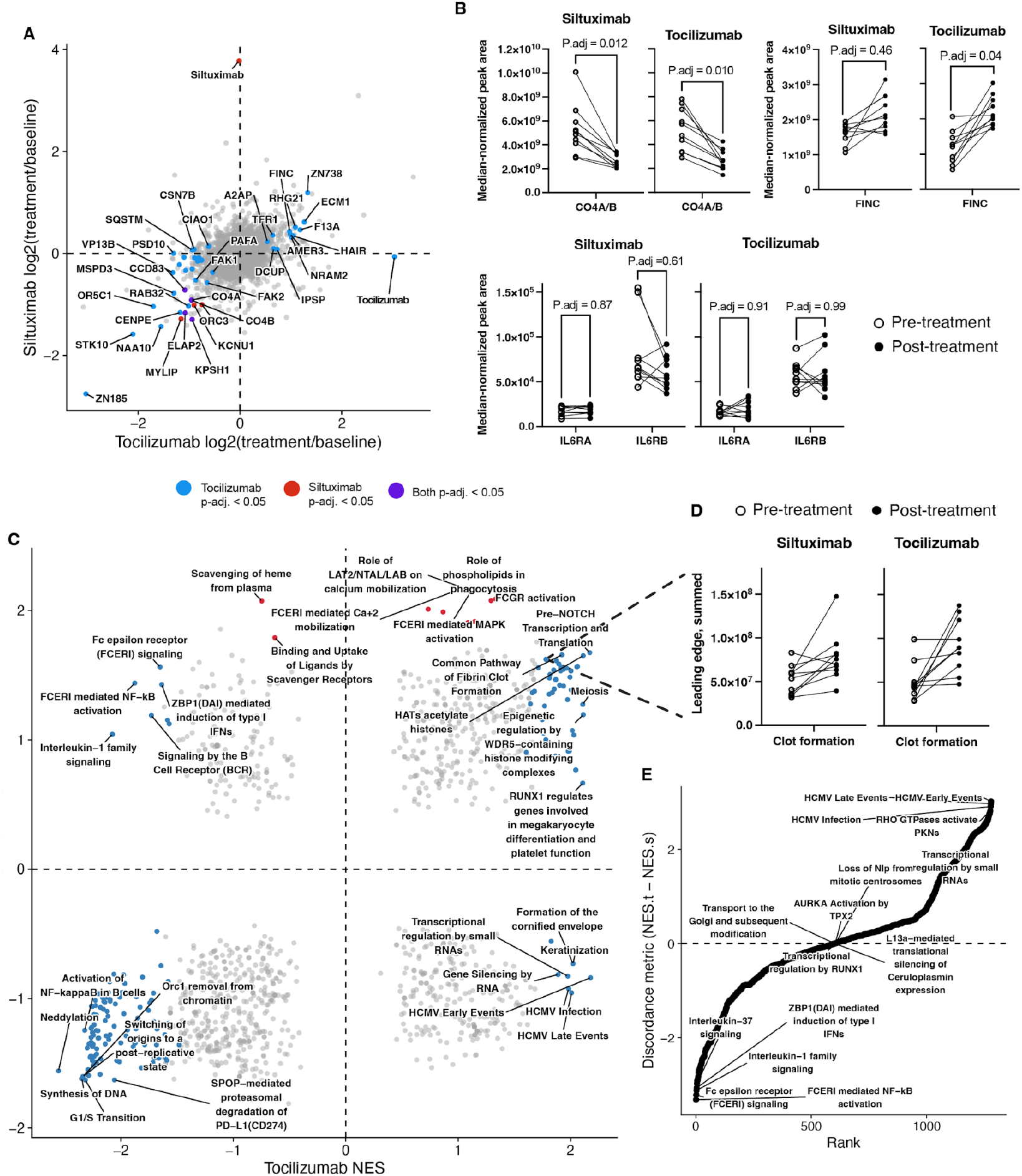
IL-6 blockade causes protein and pathway-level changes to the serum proteome. (A) Scatterplot comparison of tocilizumab and siltuximab log-fold change effects on the serum proteome. Significantly altered (BH adj. P-val < 0.05) proteins from tocilizumab shown in blue, from siltuximab shown in red, and proteins significantly altered by both treatments shown in purple. (B) Selected proteins complement factor 4 (CO4A and CO4B), IL6RA, and gp130 (IL6RB) pulled out and plotted separately. (C) genes ranked by log-2 fold change for either tocilizumab (x-axis) or siltuximab (y-axis) were evaluated for reactome pathway enrichment by fgsea. Blue = padj. for tocilizumab < 0.05, red = padj. for silxutimab < 0.05. (D) Gsea leading edge members, summed for clot formation pathway (FGB, F13A1, FGG, SERPINA5, FGA, PRTN3). (E) Discordance analysis of gsea pathways (tociliziumab NES - siltuximab NES).

IL-6 is a direct ligand of IL-6RA, with gp130 (IL-6RB) acting as a signaling transducer for the receptor complex (Figure 1A). The serum abundance of neither IL-6RA nor IL-6RB was altered in abundance by siltuximab or tocilizumab (Figure 3B). This is in agreement with our previous study, which showed that neither siltuximab nor tocilizumab altered the mbIL-6R content of CD4+ T-cells and that siltuximab did not alter either the concentration of soluble IL-6R or the cellular concentration of IL6RB^15^.

To examine the pathways altered by tocilizumab or siltuximab treatment, gene set enrichment analysis (GSEA) was performed. Fast GSEA (FGSEA)^26^ was used with the reactome database^21^ as the search space, and the log_2_ fold-change of each treatment effect as the ranking. Tocilizumab impacted a number of pathways, including a systematic decrease of neddylation, proteasome function, and interleukin 1 signaling. In contrast, siltuximab altered many fewer pathways, but among these was an increase in pathways related to Fc-gamma receptor activation, and classical complement activation. Notably, no pathway evaluated was significantly altered by both treatments (Figure 3C). Measuring the discordance as a difference in normalized enrichment scores between the two treatments (tocilizumab - siltuximab) shows FC-epsilon receptor signaling and HCMV infection pathways as behaving very differently between the two treatments. (Figure 3E).

IL-6 is known to upregulate the production of fibrinogen^12^, and inhibition of IL-6R by tocilizumab is known to reduce hypercoagulability in the context of COVID-19 infection^27^. Our data demonstrate that IL-6R blockade leads to an increase in the abundance of proteins belonging to the fibrin clot formation pathway (Figure 3D). Serum is the soluble fraction of proteins after artifactual clot induction which we expect to drive an artifactual reduction in soluble fibrinogen. Treatment with tocilizumab lead to an increase in soluble, serum-derived fibrin clot formation pathway members which suggests that IL-6R blockade is inhibiting clot formation.

### Impacts of IL-6 blockade correlate with monoclonal antibody quantitation

The serum half-life of the monoclonal antibody treatments siltuximab and tocilizumab are 21 and 11 days, respectively – long enough to quantify each antibody in the serum proteome (Figure 1 E-G). Proteomic quantitation of these therapies found individual-level variability in post-treatment serum concentration (Figure 2, C-D). Leveraging the ability to quantify the antibody therapy in the same assay as the affected serum proteome, correlational analyses were run to determine most proximal effects of therapeutic antibody treatment. For each quantified extracellular vesicle protein within a treatment cohort, the Pearson correlation coefficient was calculated as the vector of siltuximab (Figure 4A) or tocilizumab (Figure 4B) vs. each protein. Statistical significance demonstrates an association between antibody concentration and treatment effect on the extracellular vesicle proteome.

**Figure 4.**
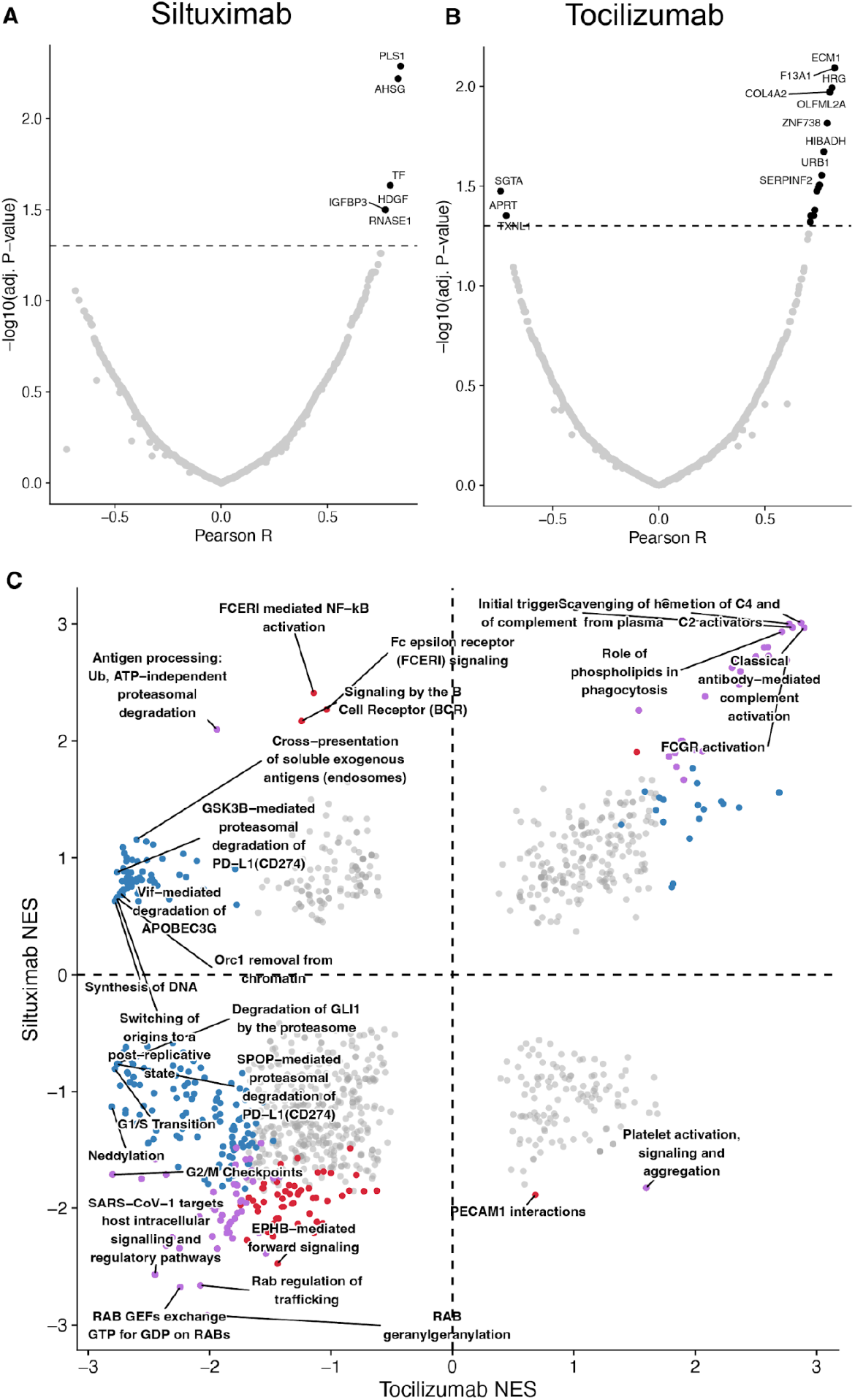
Therapeutic antibodies correlate with impacted proteins. LC-MS quantified monoclonal antibody treatments siltuximab (A) and tocilizumab (B) correlated to each quantified therapeutic antibody. (C) fgsea analysis against Reactome database, ranked by pearson R from A and B.

To determine which pathways were most represented and more correlated to antibody therapy, proteins were ranked by Pearson R and used as input for GSEA with the reactome dataset. In contrast to ranking by log_2_(fold change treatment/baseline) (Figure 3C), ranking by correlation to therapeutic antibody concentration found many pathways correlated with both IL-6 blockade (siltuximab) and IL-6R blockade (tocilizumab) (Figure 4C), including classical antibody-mediated complement activation. When GSEA is performed with the Jensen’s compartment dataset, affected proteins lie in blood microparticle and vesicle lumen compartments, consistent with the fact that we are capturing an extracellular-vesicle enriched serum matrix (Supplemental figure 1).

## Discussion

This study provides a proteomic characterization of serum responses to IL-6 pathway blockade comparing siltuximab (anti-IL-6) and tocilizumab (anti-IL-6R) in people with type 1 diabetes. Through analysis of 40 longitudinal serum samples using Mag-Net enrichment and DIA-MS, we quantified 3319 protein groups and identified 46 proteins with significantly altered abundance following treatment. While these two monoclonal antibodies produced divergent effects on T cell fate and function, our proteomics analysis reveals predominantly similar directional changes in the serum proteome, with tocilizumab showing a stronger effect (42 altered proteins) compared to siltuximab (9 altered proteins).

While we measured many individual proteins with statistically changing serum abundance with mAb treatment, a fundamental limitation in plasma proteomics remains the detection of low-abundance signaling molecules, particularly cytokines and chemokines. Despite the expanded dynamic range provided by the Mag-Net EV isolation, we were unable to directly detect IL-6 itself and other cytokines. This limited ability to detect low abundance cytokines is a well recognized challenge in mass spectrometry based proteomics, as these molecules typically circulate at pg/mL concentrations, >10^10^ lower abundance than other proteins in the serum proteome.^11^ However, borrowing lessons from genomics where pathway-level analysis has proven more robust and interpretable than individual gene changes, we can observe clear signatures of IL-6 pathway modulation through downstream effectors. These systematic alterations in acute phase proteins, complement factors, and extracellular matrix proteins provide a readable signature of IL-6 activity suppression. This pathway-level perspective is particularly valuable as it integrates information across multiple proteins, providing more statistical power and biological interpretability than individual measurements.

Several pathways appeared to differ between treatments in the fold-change–ranked GSEA, with tocilizumab showing associations with suppression of interleukin-1 and B-cell receptor signaling, while siltuximab effects were limited to Fc-gamma receptor activation and classical complement pathways. However, given the markedly greater number of proteins reaching significance with tocilizumab, these apparent divergences may largely reflect differences in statistical power or effective dose rather than fundamentally distinct downstream biology. Consistent with this interpretation, ranking proteins by their correlation to measured antibody levels (Figure 4C) revealed substantially more pathway overlap between treatments, suggesting that the core biology of IL-6 pathway blockade is shared, but detected at different sensitivities in this study. The broader proteomic footprint of tocilizumab is also consistent with the expected consequences of receptor-level versus ligand-level blockade: while siltuximab neutralizes IL-6 alone, tocilizumab blocks IL-6R and thereby may disrupt trans-signaling mediated by soluble IL-6R/gp130 complexes as well as signaling by other IL-6 family cytokines that share gp130 as a signal transducer, potentially affecting a wider range of downstream pathways. Nonetheless, further study would be needed to definitively separate potency-driven differences in signal detection from true mechanistic divergence.

This distinction is relevant for understanding IL-6 biology, as this cytokine coordinates responses across multiple organ systems including liver (acute phase response), bone marrow (hematopoiesis) and endothelium (vascular permeability)^12^. The similar serum proteome signatures may reflect the dominant contribution of hepatic acute phase responses, which could mask more subtle cell-type-specific differences observed in isolated immune populations. Indeed, both treatments showed a similar suppression of complement factors C4A/B, consistent with previous reports of IL-6 blockage reducing complement activation, likely through hepatic mechanisms.

The broadly similar response of the serum proteome to IL-6 or IL-6R inhibition represents a contrast to our previous study which demonstrated that different modes of IL-6 inhibition have differential impacts to the cellular immune response^15^. This contrast highlights an important point: therapeutic interventions can produce different effects at different scales and different sites. Our previous work focused on isolated T cell populations, providing a specific and localized view of cell-autonomous response. In contrast, our serum proteomics captures a broader systemic response across tissues and cell types.

This study demonstrates several technical advances important for clinical proteomics. The excellent correlation (Spearman ρ > 0.95) between baseline samples from the same individuals collected a year apart validates the robustness of this approach for longitudinal studies. In addition, our peptide-centric analysis of therapeutic antibodies showcases how bottom-up proteomics can distinguish highly homologous proteins within complex mixtures. By identifying unique peptides that change with treatment we could specifically track therapeutic antibodies despite extensive sequence similarity with endogenous immunoglobulins. This approach could be broadly applicable for monitoring other biological therapeutics. In addition, the ability to profile individual treatment responses in serum or plasma has important implications for precision medicine. Our observation that proteomics responses correlate with measurable drug levels suggests that therapeutic drug monitoring could be enhanced by simultaneous measurement of pharmacodynamic markers. The proteins identified as most strongly correlating with antibody levels could serve as candidate markers for treatment response or target engagement. For understanding immunomodulatory effects relevant to autoimmunity, cellular assays may continue to be most informative, but for monitoring systemic inflammation or predicting adverse events, serum proteomics may provide relevant information.

This study demonstrates that comprehensive serum proteomics can reveal systemic responses to therapeutic interventions that complement targeted cellular analyses. These findings underscore the importance of applying different assays to assess the biological response to treatment and different scales. As mass spectrometry based proteomics technology continues to advance, the ability to deeply profile individual responses will become increasingly valuable for optimizing therapeutic selection and monitoring in precision medicine.

## Notes

The authors declare the following competing financial interest(s): DLP is an employee of Thermo Fisher Scientific, the manufacturer of the mass spectrometry instrumentation used in this research. The MacCoss lab at the University of Washington has a sponsored research agreement with Thermo Fisher Scientific, the manufacturer of the mass spectrometry instrumentation used in this research. DLP is an employee of Thermo Fisher Scientific. D.K.S. is a consultant and/or collaborator with ThermoFisher Scientific, AI Proteins, Genentech, and Matchpoint Therapeutics. The other authors have no conflicts to declare.

## Acknowledgments

The authors would like to thank the MacCoss and Schweppe Labs for their helpful advice and comments. Funding and drugs for the siltuximab study were provided by Janssen Pharmaceuticals under a contract research agreement with CJG and JHB. This work was funded in part by National Institutes of Health grants U01 DK137097 and R24 GM141156. The tocilizumab study was funded by Juvenile Diabetes Research Foundation (JDRF) under a partnership with the Immune Tolerance Network (JDRF grant 2-PAR-2016-349-Q-R to GTN).

---

**Supplemental figure 1.**
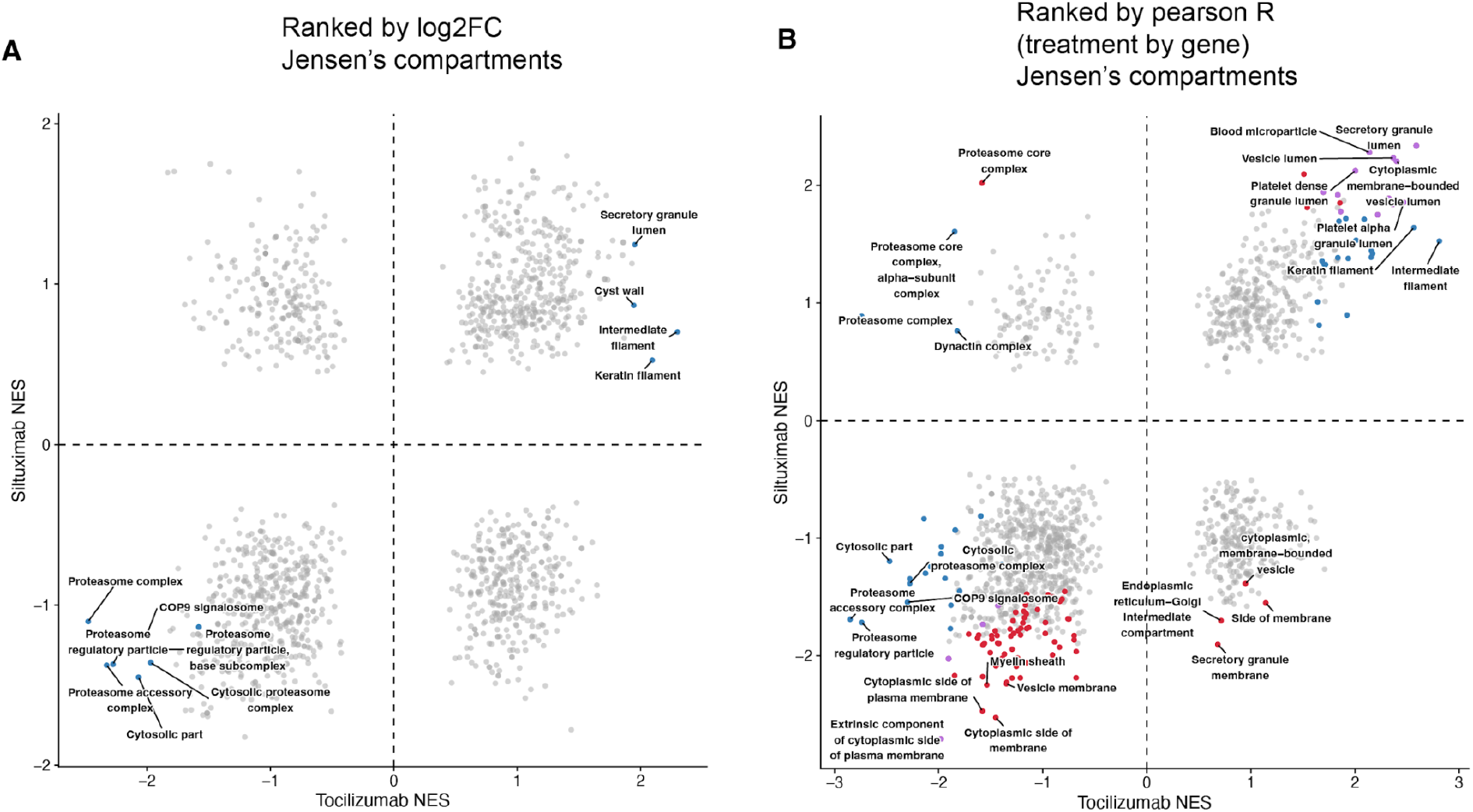
Fgsea analyses with Jensen’s compartments at the gene set. (A) Protein groups ranked by log2FC used as input. (B) Protein groups ranked by pearson R to corresponding treatment mAb (tocilizumab or siltuximab).

